# Construction and Experimental Validation of a Petri net Model of Wnt/β-catenin Signaling

**DOI:** 10.1101/044966

**Authors:** Annika Jacobsen, Nika Heijmans, Folkert Verkaar, Martine J. Smit, Jaap Heringa, Renée van Amerongen, K. Anton Feenstra

## Abstract

The Wnt/β-catenin signaling pathway is important for multiple developmental processes and tissue maintenance in adults. Consequently, deregulated signaling is involved in a range of human diseases including cancer and developmental defects. A better understanding of the intricate regulatory mechanism and effect of physiological (active) and pathophysiological (hyperactive) WNT signaling is important for predicting treatment response and developing novel therapies. The constitutively expressed CTNNB1 (commonly and hereafter referred to as β-catenin) is degraded by a destruction complex, composed of amongst other AXIN1 and GSK3. The destruction complex is inhibited during active signaling leading to β-catenin stabilization and induction of β-catenin/TCF target genes. In this study we investigated the mechanism and effect of β-catenin stabilization during active and hyperactive WNT signaling in a combined *in silico* and *in vitro* approach. We constructed a Petri net model of Wnt/β-catenin signaling including main players from the plasma membrane (WNT ligands and receptors), cytoplasmic effectors and the downstream negative feedback target gene *AXIN2*. We simulated the model with active (i.e. WNT stimulation) and hyperactive (i.e. GSK3 inhibition) signaling, which led to the following observations: 1) A dose- and time-dependent response was observed for both WNT stimulation and GSK3 inhibition. 2) The Wnt-pathway activity was 2-fold higher for GSK3 inhibition compared to WNT stimulation. Both of these observations were corroborated by TCF/LEF luciferase reporter assays. Using this experimentally validated model we simulated the effect of the negative feedback regulator AXIN2 upon WNT stimulation and observed an attenuated β-catenin stabilization. We furthermore simulated the effect of APC inactivating mutations, yielding a stabilization of β-catenin levels comparable to the Wnt-pathway activities observed in colorectal and breast cancer. Our model can be used for further investigation and viable predictions of the role of Wnt/β-catenin signaling in oncogenesis and development.

**Author Summary:** Deregulated Wnt/β-catenin signaling is implicated in cancer and developmental defects. In this study we combined *in silico* and *in vitro* efforts to investigate the behavior of physiological and pathophysiological WNT signaling. We created a model of Wnt/β-catenin signaling that describes the core interactions: receptor activation, inhibition of downstream effectors and an important negative feedback mechanism. Simulations with the model demonstrated the expected dose- and time-dependent response for both conditions, and the Wnt-pathway activity was significantly higher for pathophysiological compared to physiological signaling. These observations were experimentally validated, which allowed us to investigate and predict the effect of the negative feedback and an inactivating cancer mutation on the Wnt-pathway activity. Our model provides mechanistic insight on the different conditions and can easily be extended and used to answer other questions on Wnt/β-catenin signaling in the area of cancer research and regenerative medicine.

## Introduction

The Wnt/β-catenin signaling pathway is crucial for regulating cell proliferation and differentiation during embryonic development, while in adults it helps control tissue homeostasis and injury repair in stem cell maintenance [1, 2]. Extracellular WNT ligands activate signaling leading to CTNNB1 (commonly and hereafter referred to as β-catenin) stabilization, nuclear translocation, interaction with TCF/LEF transcription factors [3] and induction of β-catenin/TCF target genes [4] (Fig 1B). A critical feature of Wnt/β-catenin signaling is the inhibition of a ‘destruction complex’ which degrades the constitutively expressed β-catenin (Fig 1A) [5].

**Fig 1.**
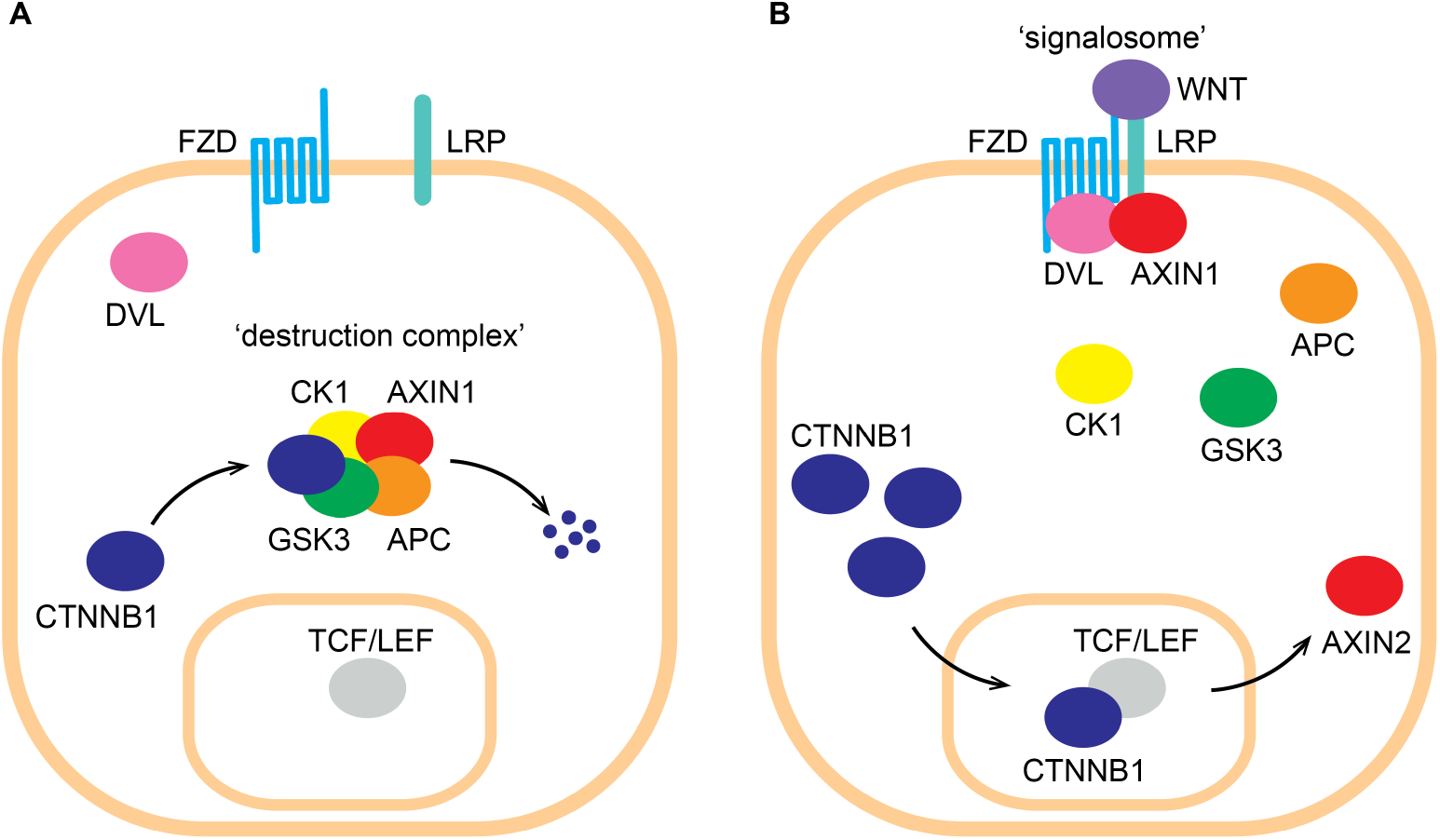
Illustration of Wnt/β-catenin signaling. (A) In the absence of an external WNT stimulus β-catenin (referred to by its official gene name CTNNB1 in the figure) is continuously degraded by a ‘destruction complex’ consisting of AXIN1, adenomatous polyposis coli (APC), casein kinase 1 (CK1) and glycogen synthase kinase 3 (GSK3). (B) Extracellular WNT interacts with the membrane-bound receptors frizzled (FZD) and lipoprotein receptor-related protein (LRP). Dishevelled (DVL) interacts with the intracellular tail of FZD and sequesters AXIN1 to the plasma membrane forming a so-called ‘signalosome’. The ensuing depletion of the cytoplasmic pool of AXIN1 inhibits the formation of the destruction complex. β-catenin thereby stabilizes and translocates to the nucleus, where it interacts with TCF/LEF transcription factors activating transcription of specific target genes including *AXIN2*.

The destruction complex consists of two scaffolding proteins, AXIN1 and adenomatous polyposis coli (APC), and two kinases, casein kinase 1 (CK1) and glycogen synthase kinase 3 (GSK3). β-catenin is phosphorylated by CK1 and GSK3 [6, 7] and thereafter presented to the proteasome for ubiquitination [8] and degradation (Fig 1A). Extracellular WNT binds to and activates the 7 transmembrane receptor, Frizzled (FZD) [9], and the co-receptor, lipoprotein receptor-related protein (LRP5/6) [10]. The intracellular tail of FZD interacts with Dishevelled (DVL) through an incompletely understood mechanism and sequesters AXIN1 to the cell membrane [11] forming a so-called ‘signalosome’ [12]. This leads to depletion of the cytoplasmic pool of the destruction complex component AXIN1, which in turn inhibits the formation of the destruction complex itself (Fig 1B). It is not fully understood whether only AXIN1 or more destruction complex components are sequestered to the cell membrane during WNT signaling. Indeed, a study by Li *et al*. [13] showed that AXIN1 does not dissociate from the other destruction complex components during WNT signaling.

The inhibition of the destruction complex leads to β-catenin stabilization and nuclear translocation. Nuclear β-catenin interacts with TCF/LEF transcription factors [14] forming the β-catenin/TCF transcriptional (co)activator complex. A collection of more than 100 genes induced by β-catenin/TCF transcription is listed on the WNT homepage (www.web.stanford.edu/group/nusselab/cgi-bin/wnt/) (last update: September 2015). The specific subset of genes induced, however, strongly depends on tissue type and developmental stage [15]. Several of these target genes are feedback regulators, where *AXIN2* is of particular interest. First, *AXIN2* is a universal β-catenin/TCF target gene and as such it is believed to faithfully report Wnt-pathway activity in multiple tissues [16, 17]. Second, AXIN2 encodes a functional homolog of the destruction complex component AXIN1 [18] and mediates an auto-inhibitory feedback loop. Although AXIN1 and AXIN2 share functional similarities, they are only partially redundant *in vivo* due to their different expression patterns [19]: *AXIN1* is constitutively expressed [20], whereas *AXIN2* is induced during active Wnt/β-catenin signaling [17, 21]. The AXIN2 negative feedback is believed to be important for the tight spatio-temporal regulation of Wnt/β-catenin signaling [22]. However, the exact regulatory role of AXIN2 remains an open question.

Deregulated Wnt/β-catenin signaling caused by genetic alterations can have major developmental consequences, and is the leading cause of colorectal oncogenesis [23]. The most common colorectal cancer mutation is found in APC [24, 25]. Different APC inactivating mutations lead to different levels of Wnt-pathway activity e.g. higher β-catenin stabilization is seen in colorectal cancer compared to breast cancer (as reviewed in [26]). Other rarer colorectal cancer mutations [27] are found in AXIN1 [28], AXIN2 [29, 30] and β-catenin [31, 32]. As a common mode of action, these oncogenic mutations cause hyperactive WNT signaling [33].

Investigating the mechanism and effect of β-catenin stabilization during physiological (active) and pathophysiological (hyperactive) WNT signaling is crucial for developing effective treatment, both in the field of cancer research and regenerative medicine. *In vitro* experiments in which cells are stimulated with WNT are generally assumed to represent active signaling, whereas downstream oncogenic mutations represent hyperactive signaling. Inhibition of GSK3 using small molecule inhibitors is widely used to activate WNT signaling during cellular reprogramming and in embryonic stem cell cultures [34, 35]. Inhibition of GSK3 inhibits the destruction complex, which can be interpreted as similar to the effects of oncogenic mutations. Several mathematical models of Wnt/β-catenin signaling have been created as reviewed in [36] to facilitate these investigations. However the construction of these models requires detailed information on e.g. protein concentrations and reaction rates, which require large experimental efforts. Consequently, the currently available models include many estimated parameters, which limits their scale of applicability [36]. On the other hand, coarse-grained data on interactions and relative levels of proteins are readily available, and much easier to obtain. Enabling the use of such data would greatly expand the scale of applicability of modeling. With this in mind, we previously introduced a Petri net modeling formalism that can utilize this type of coarse-grained data [37, 38].

In this paper, we present a combined computational and experimental approach to build on the investigations of the mechanism and effect of β-catenin stabilization during active and hyperactive WNT signaling. We created a Petri net model of Wnt/β-catenin signaling describing membrane activation by the WNT ligand, β-catenin degradation by the destruction complex and the negative feedback by AXIN2. We used the model to explain how active signaling upon WNT stimulation and hyperactive signaling upon GSK3 inhibition leads to different levels of β-catenin stabilization. We corroborated our observations from the model using data from TCF/LEF luciferase reporter assays and Western blot analysis. We then used the experimentally validated model to explore plausible modes of action of β-catenin stabilization as a result of negative feedback by activating expression of AXIN2 upon WNT stimulation, or due to APC inactivating mutations that are known to play a key role in oncogenesis of colorectal and breast cancer.

## Results

### Modeling Wnt/β-catenin signaling

We created a Petri net model of Wnt/β-catenin signaling to investigate the mechanism and effect of β-catenin stabilization under physiological (e.g. embryonic development) and pathophysiological (e.g. cancer) conditions (Fig 2). The model describes the interactions between the core proteins in the pathway with a focus on capturing the behavior of β-catenin stabilization following destruction complex inhibition. Therefore the degradation of β-catenin by the destruction complex is specifically in the model. Likewise, the gene expressions of *β-catenin* (encoding β-catenin), but also of *AXIN2*, a negative feedback target gene of the Wnt-pathway, are also specifically included in the model. Production and degradation of all other proteins are assumed to have similar rates and are therefore omitted, such that the token levels of these proteins remain the same throughout the simulation (See Materials and Methods).

**Fig 2.**
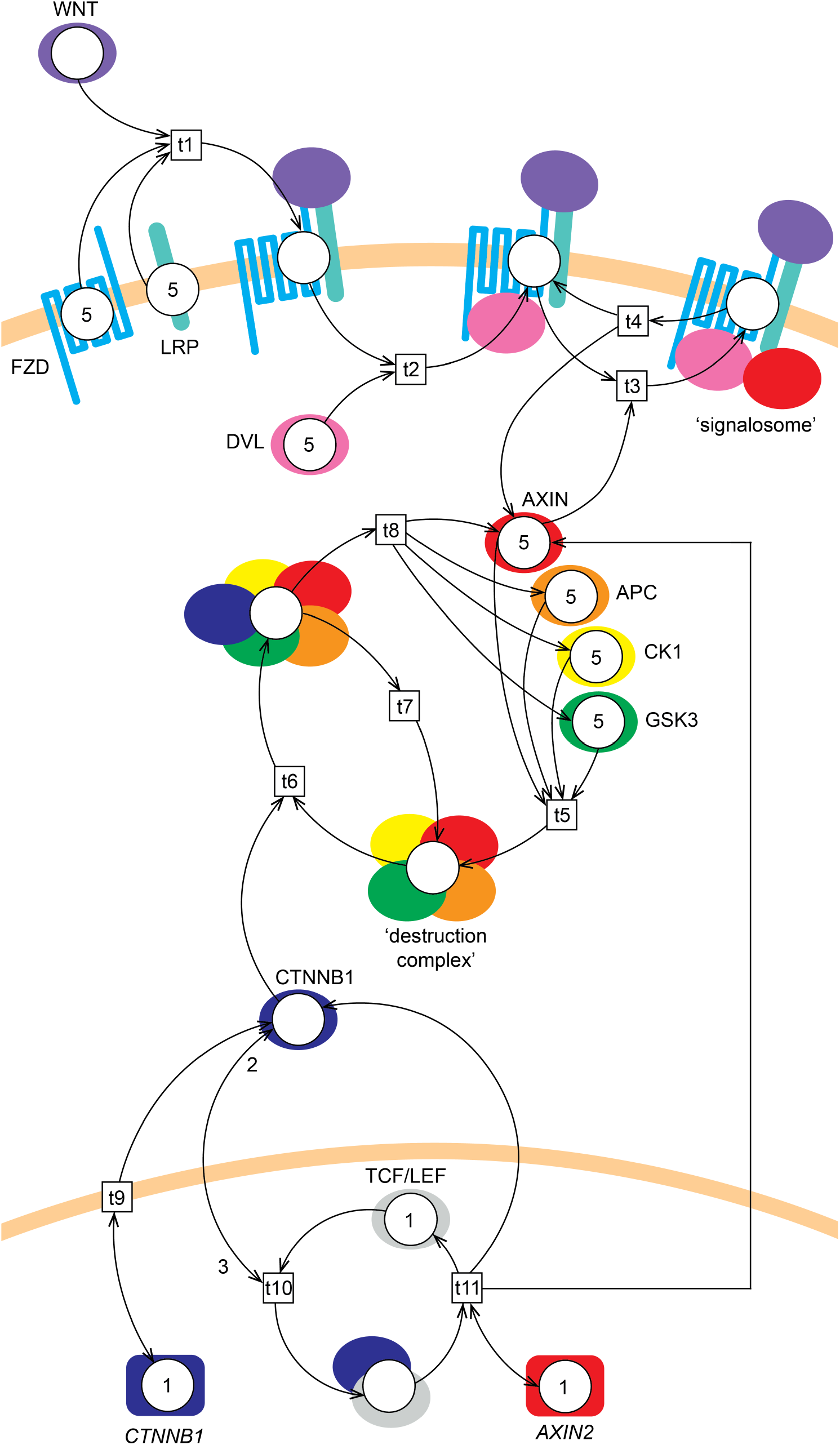
Petri net model of Wnt/β-catenin signaling. The model consists of 18 places (circles, representing gene or protein states), 11 transitions (boxes, representing protein complex formation, dissociation, translocation or gene expression) and 40 arcs (arrows, representing the direction of flow of the tokens). WNT initiates signaling by binding to FZD and LRP (t1), forming the WNT/FZD/LRP complex. DVL and AXIN1 then interact with this complex intracellularly (t2 and t3, respectively) forming a so-called ‘signalosome’. The signalosome dissociates once every 10 steps (t4) into WNT/FZD/LRP/DVL and AXIN1. β-catenin is referred to by its official gene name CTNNB1 in the figure. The β-catenin protein is produced every step (t9) by the *β-catenin* gene. AXIN1, APC, CK1 and GSK3 interact (t5) and form a ‘destruction complex’. The destruction complex binds β-catenin (t6) to mark it for degradation. The destruction complex is then either reused (t7) for another round of β-catenin degradation or dissociates (t8) into its components AXIN1, APC, CK1 and GSK3. Alternatively, β-catenin can interact with TCF/LEF in the nucleus (t10), leading to transcriptional activation of *AXIN2* (t11). Initial token levels are 0 (not shown), 1 or 5 (depicted in the places). Most arc weights are 1 (not shown), except for the nuclear translocation and interaction of β-catenin to TCF/LEF transcription factors, which has an incoming arc weight of 3 and an outgoing arc weight of 2 (depicted on the arcs).

The model consists of 18 places (circles, representing gene or protein states), 11 transitions (boxes, representing protein complex formation, dissociation, translocation or gene expression) and 40 arcs (arrows, representing the direction of flow of the tokens). In the model, WNT initiates signaling extracellularly by binding to its transmembrane receptors FZD and LRP (t1), forming the WNT/FZD/LRP complex. DVL interacts with the intracellular tail of FZD when present in the WNT/FZD/LRP complex (t2), forming the WNT/FZD/LRP/DVL complex. DVL thereafter sequesters AXIN1 to the membrane (t3) forming the signalosome consisting of WNT, FZD, LRP, DVL and AXIN1. In the model we have not included the contribution of GSK3 and CK1 in the formation of the signalosome, because these two multitasking kinases are generally assumed not to be rate-limiting in the cell [23, 39]. Further, AXIN1 is the only destruction complex constituent that binds to the signalosome in the model. The signalosome dissociates once every 10 steps (t4) into the WNT/FZD/LRP/DVL complex and AXIN1 in order to incorporate a lower dissociation- than formation-rate of the signalosome. The destruction complex, which sequesters β-catenin unless WNT induces signalosome formation, is formed (t5) by AXIN1, APC, CK1 and GSK3. In the model, β-catenin binding to the destruction complex leads to degradation of β-catenin (t8 and t7), and the destruction complex is then either reused (t7) for another round of β-catenin degradation or dissociates (t8) to AXIN1, APC, CK1 and GSK3. In the model, β-catenin protein is produced every step (t9) following transcription of the *β-catenin* gene, and either binds the destruction complex (t6) or translocates to the nucleus, where it interacts with TCF/LEF (t10) to activate transcription of *AXIN2* (t11). Since AXIN1 and AXIN2 are functional homologs [18], they are modeled as one protein entity (depicted as ‘AXIN’). Further, we do not distinguish between the cytoplasmic and nuclear pool of β-catenin in the model. This allowed the nuclear translocation and TCF/LEF interactions to be modeled as one transition (t10).

The initial token level of all protein places was set to 5, except for TCF/LEF, which was set to 1, and β-catenin, which was set to 0. The initial token level of all protein complexes was set to 0. The initial token level of the gene places, *β-catenin* and *AXIN2*, was set to 1 (since these genes are always presumed to be present). Most arc weights were set to 1, with an exception of the arc weight from β-catenin to transition t10 (i.e. its translocation to the nucleus and subsequent interaction with TCF/LEF), which was set to 3, and the arc weight from t10 to β-catenin, which was set to 2. From the model point of view this means that for t10 to fire, the β-catenin place needs a level of 3 tokens, but that only 1 is consumed (See Fig 2). These weights were chosen because it is generally believed that β-catenin accumulates in the cytoplasm before it translocates to the nucleus and binds TCF/LEF. Parts of this initial setup were changed accordingly to mimic the different conditions of Wnt/β-catenin signaling simulated in this study (see below).

### Active signaling upon WNT stimulation

We simulated WNT stimulation to predict the level of β-catenin stabilization during active signaling. To this end we ran a series of simulations with different initial WNT token levels (ranging from 0 to 5) without AXIN2 feedback (i.e. the arc weight from t11 to AXIN was set to 0). As shown in Fig 3A, we observed four different β-catenin response levels depending on the initial WNT token level. A flat β-catenin response was seen for WNT = 0, 1 or 2. For WNT = 3, 4 or 5, we observed a delay in the initial increase of β-catenin, which eventually increased linearly with a slope depending on the WNT level. The β-catenin stabilization was low for WNT = 3 and moderate for WNT = 4 and 5. Maximal WNT stimulation (WNT = 5) led to a stabilization of ~60 β-catenin tokens.

High-throughput analyses of Wnt-pathway activation (i.e. a comparison of multiple doses and time points within the same experiment) can be performed using a TCF/LEF luciferase reporter assay, which faithfully reports Wnt/β-catenin signaling [40]. To validate the β-catenin levels predicted upon WNT stimulation by our model, we treated HEK293T^WOO^ cells (carrying a stably integrated β-catenin/TCF luciferase reporter) with increasing concentrations of purified, commercially available, Wnt3a for 3, 8 and 24 hours. These experiments reproduce the dose- and time-dependent increase of TCF/LEF reporter gene activity predicted above, thereby validating our model (Fig 3B and 3C). To directly link the results from the reporter gene assay to an increase in β-catenin protein levels, we repeated the experiment for one level of WNT stimulation (100 ng/ml purified Wnt3a) for a more extensive time series, including additional earlier time points, and analyzed the results by performing both a TCF/LEF reporter gene assay (Fig 3B and 3D) and quantitative Western blot analysis (Fig 3E and 3F). The latter allows direct, albeit less sensitive, detection of β-catenin protein levels. Both the transcriptional reporter assay and the measurement of β-catenin protein levels show a time-dependent increase (Fig 3D-3F). Direct comparison of the two readouts reveals the inherent limitations of each of the two experimental systems: The change (i.e. fold increase) in TCF/LEF reporter activity is more pronounced than, but slightly delayed compared to, the change in β-catenin protein levels.

**Fig 3.**
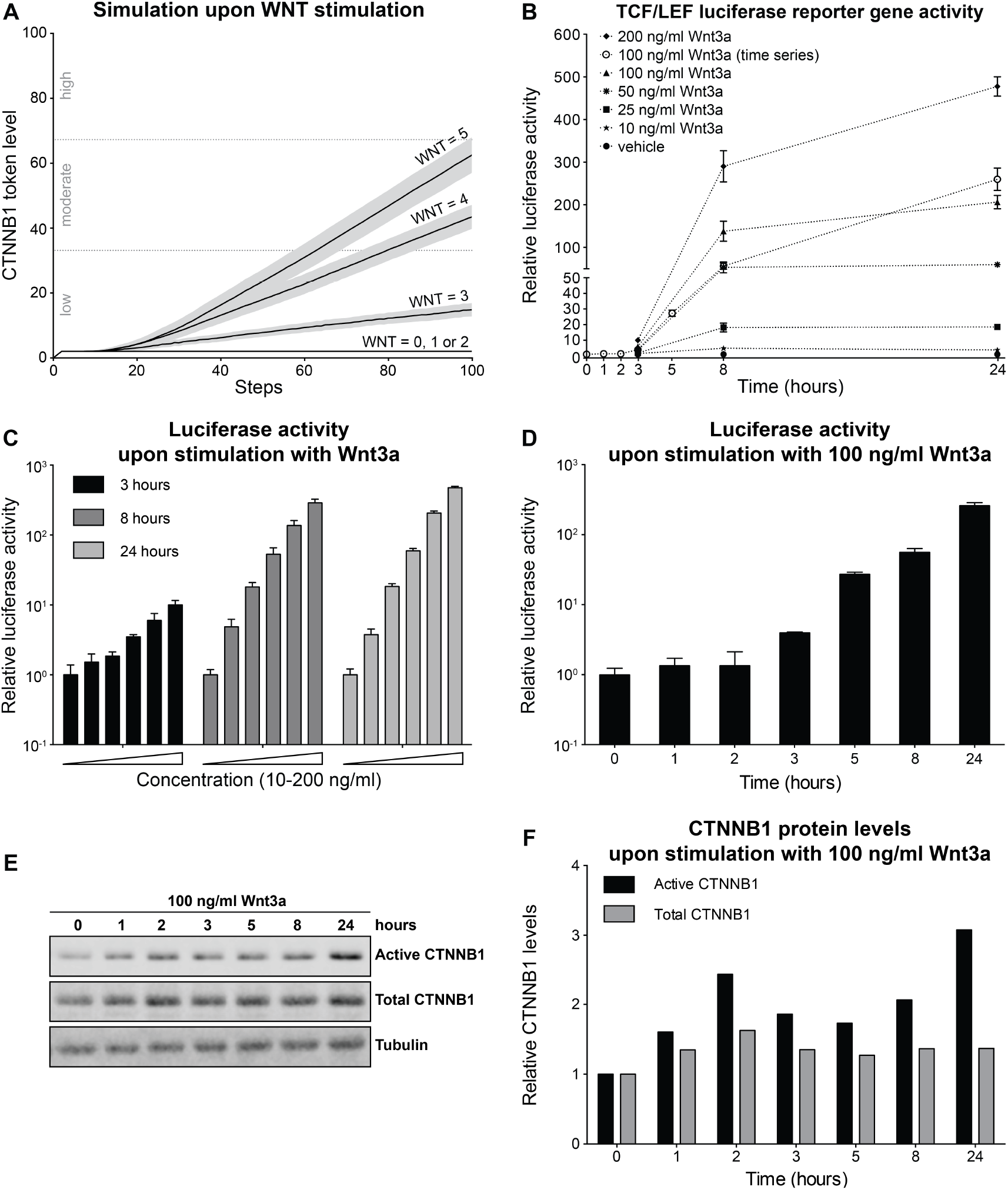
Model simulation and experimental validation of Wnt-pathway activation upon WNT stimulation. (A) β-catenin (referred to by its official gene name CTNNB1 in the figure) token levels predicted by our model with initial WNT token levels ranging from 0 to 5. For WNT = 0, 1 or 2, we observed a flat β-catenin response. For WNT = 3, 4 and 5 β-catenin increases from low to moderate levels. (B) Reporter assay in HEK293T^WOO^ cells, showing dose- and time-dependent activation of a Wnt/β-catenin responsive TCF/LEF luciferase reporter to allow easy comparison to the model results in panel (A). For all conditions shown in black (corresponding to panel C), luciferase activity was plotted relative to the vehicle control (not shown), which was set at 1 for each of the three time points (3, 8 and 24 hours). For the curve shown in white (corresponding to panel D), luciferase activity was plotted relative to the vehicle control, which was set at 1 for the t=0 hours condition. (C) Reporter assay in HEK293T^WOO^ cells, showing dose-dependent activation at 3, 8 and 24 hours after stimulation with purified Wnt3a (same data as in B). (D) Reporter assay in HEK293T^WOO^ cells, showing time-dependent activation upon treatment with 100 ng/ml of Wnt3a. Values were plotted relative to the vehicle control, which was set at 1 for t=0 hours. (E) Western blot analysis from the experiment depicted in (D), showing total and active (non-phosphorylated) β-catenin levels. Tubulin was used as a loading control. (F) Quantification of the Western blot shown in (E). Total and active β-catenin levels were normalized to tubulin. The increase in either total or active β-catenin levels was plotted relative to time point 0, for which the normalized levels were set to 1.

### Hyperactive signaling upon GSK3 inhibition

To predict the level of β-catenin stabilization during hyperactive signaling by a downstream perturbation, we next simulated our model upon GSK3 inhibition. We ran a series of simulations with different initial GSK3 token levels (ranging from 5 to 0), where 5 initial tokens represents wildtype (i.e. no Wnt-pathway activity) and 0 corresponds to complete inhibition (hyperactive signaling). The simulations revealed that the response levels depend on initial GSK3 token levels (see Fig 4A). For GSK3 = 3, 4 or 5, we observed a flat β-catenin response. A linear increase in β-catenin levels with a slope depending on GSK3 levels was seen for GSK3 = 0, 1 or 2. This corresponds to β-catenin degradation ranging from no degradation to 1 or 2 β-catenin tokens degraded per three simulation steps, respectively. Consequently, β-catenin stabilization was low for GSK3 = 2, moderate for GSK3 = 1 and high for GSK3 = 0. Complete GSK3 inhibition led to a stabilization of 100 β-catenin tokens.

To validate the coarse-grained β-catenin levels predicted by our model upon GSK3 inhibition, we stimulated HEK293T^WOO^ cells with increasing concentrations of CHIR99021, one of the most potent and selective GSK3 inhibitors available to date, over a broad time range (3, 8 and 24 hours). The measured TCF/LEF reporter gene activity confirmed the dose- and time-dependent increase upon GSK3 inhibition (Fig 4B and 4C) predicted by our model (Fig 4A). As with the Wnt3a treatment, here we also performed a TCF/LEF reporter gene assay and quantitative Western blot analysis side by side for one of the treatment conditions (3 µM CHIR99021) for multiple time points. An increase in both active (i.e. non-phosphorylated) and total (i.e. both phosphorylated and non-phosphorylated) β-catenin is apparent after 1 hour, whereas an increase in the signal of the luciferase reporter assay can only be detected after 3 hours. Furthermore, the dynamic range of the Western blot analysis is limited compared to the reporter gene assay, allowing us to measure at most a 4-fold increase in β-catenin levels in the former, but up to a 10^4^ fold increase in Wnt-pathway activity in the latter (Fig 4D-4F).

**Fig 4.**
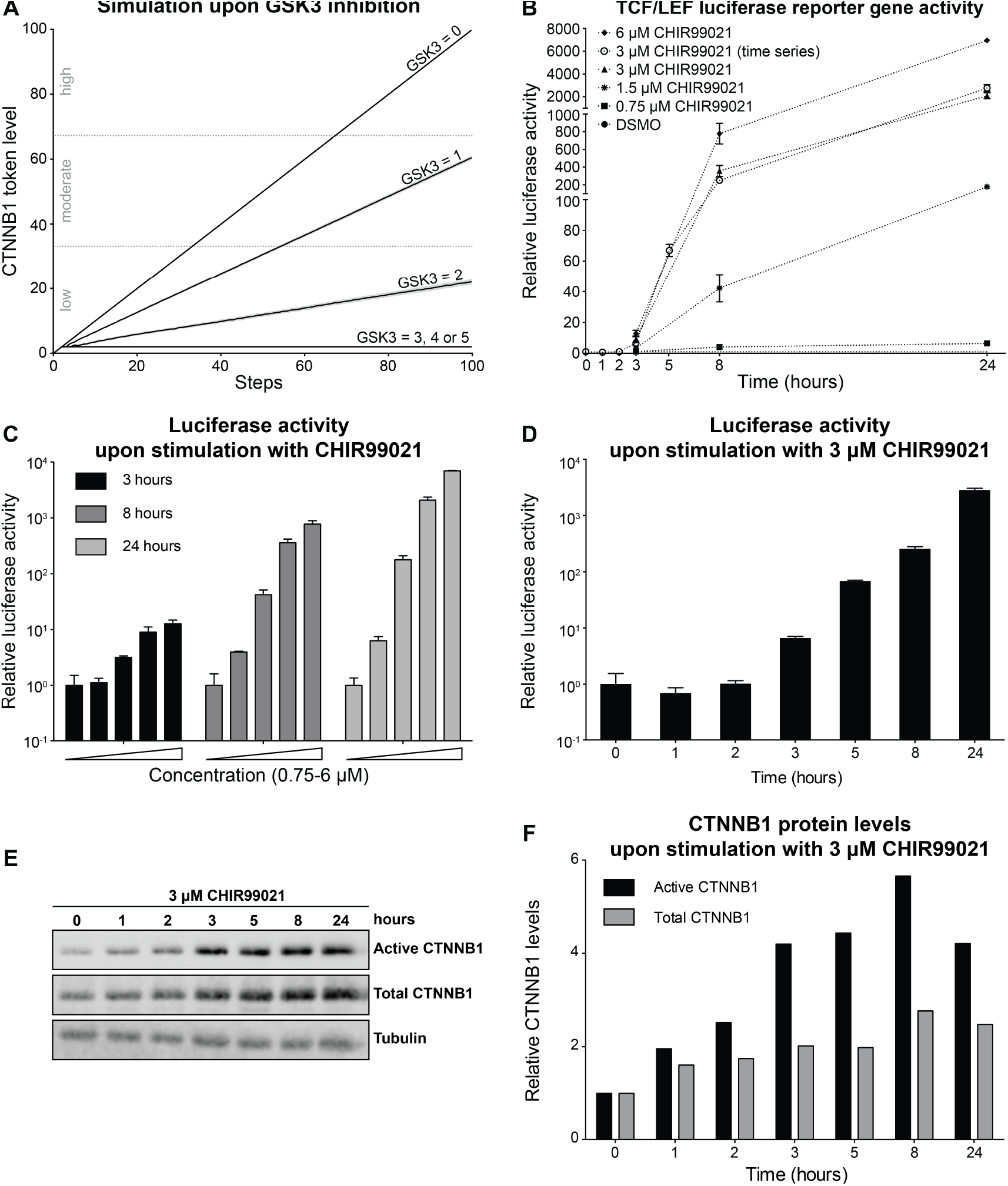
Model simulation and experimental validation of Wnt-pathway activation upon GSK3 inhibition. (A) β-catenin (referred to by its official gene name CTNNB1 in the figure) token levels predicted by our model with initial GSK3 token levels ranging from 0 to 5. For GSK3 = 3, 4 or 5, we observed a flat β-catenin response. For GSK3 = 0, 1 or 2 β-catenin increases to low, moderate or high levels, respectively. (B) Reporter assay in HEK293T^WOO^ cells, showing dose- and time-dependent activation of a Wnt/β-catenin responsive TCF/LEF luciferase reporter to allow easy comparison to the model results in panel (A). For all conditions shown in black (corresponding to panel C), luciferase activity was plotted relative to the vehicle control (not shown), which was set at 1 for each of the three time points (3, 8 and 24 hours). For the curve shown in white (corresponding to panel D), luciferase activity was plotted relative to the vehicle control, which was set at 1 for the t=0 hours condition. (C) Reporter assay in HEK293T^WOO^ cells, showing dose-dependent activation at 3, 8 and 24 hours after stimulation with CHIR99021 (same concentrations as depicted in B). (D) Reporter assay in HEK293T^WOO^ cells, showing time-dependent activation upon treatment with 3 mM CHIR99021. Values were plotted relative to the DMSO control, which was set at 1 for t=0 hours. (E) Western blot analysis from the experiment depicted in (D), showing total and active (non-phosphorylated) β-catenin levels. Tubulin was used as a loading control. (F) Quantification of the Western blot shown in (E). Total and active β-catenin levels were normalized to tubulin. The increase in either total or active β-catenin levels was plotted relative to time point 0, for which the normalized levels were set to 1.

### Predictions of hyperactive signaling by APC inactivating mutations

The most common colorectal oncogene, *APC*, perturbs downstream WNT signaling. Different APC mutations exist that result in truncated proteins negatively influence the formation of the destruction complex to different degrees. As a result, the different APC mutations lead to different levels of β-catenin stabilizations. According to a recent review [26], the β-catenin signaling activity (β-catenin reporter activity) was between 10-20% for APC mutations in breast tumors, versus 20-100% in colorectal tumors.

We used our validated model to explore if the effect of these APC mutations might be explained by different rates of destruction complex formation. We implemented the effect of the APC mutations, by decreasing the rate of the destruction complex formation, ranging from no production at all to production every 20, 10 and 5 steps. In Fig 5 we observed four different response levels for the different APC mutations, where stabilization of β-catenin levels went from low to high depending on this rate of destruction complex formation. Comparing these token levels to the β-catenin signaling activities reviewed in [26], the three highest β-catenin stabilizations would correspond to hyperactive signaling by APC mutations in colorectal tumor formation, whereas the lowest β-catenin stabilization would correspond to the effects by APC mutations as observed in breast tumor formation.

**Fig 5.**
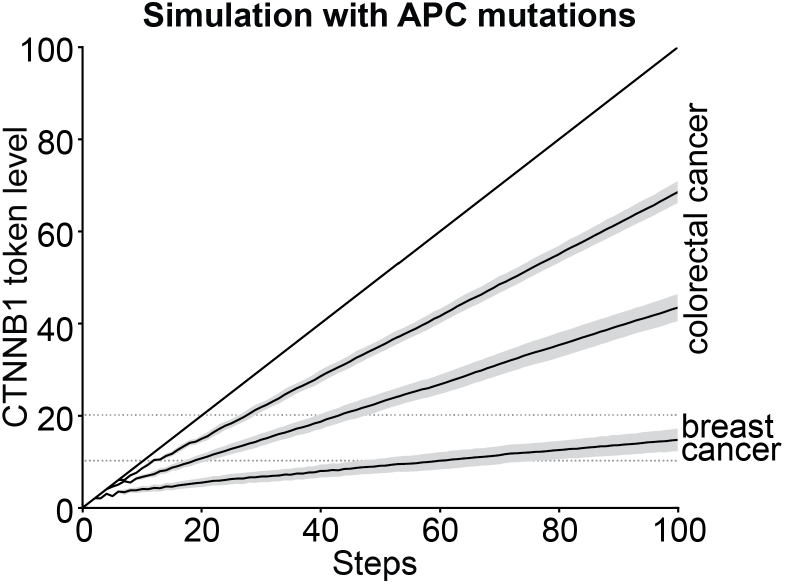
Model prediction of Wnt-pathway hyperactivation by APC inactivating mutations. β-catenin (referred to by its official gene name CTNNB1 in the figure) token levels predicted by our model with four different APC mutations (ranging from no production to production every 20, 10 and 5 steps). The highest β-catenin stabilizations might correspond to the effects by mutations in colorectal tumor formation and the lowest β-catenin stabilizations might correspond to the effects by mutations in breast tumor formation.

### Predictions of active signaling upon WNT stimulation with AXIN2 feedback

In our model *AXIN2* is induced by β-catenin/TCF transcription and increases the cytoplasmic pool of AXIN, which under certain conditions, e.g. WNT stimulation, is the limiting factor for β-catenin degradation. However, our experimental dataset obtained using Wnt3a stimulation showed no obvious decrease in β-catenin levels that might be due to this negative feedback (Fig 3E and 3F). It should be noted that in this experimental setting (100 ng/ml Wnt3a), the Wnt-pathway is likely still activated at supra-physiological levels. Moreover, the WNT ligand remains present throughout the experiment. *In vivo*, however, physiological Wnt-pathway activation is strictly regulated both due to the WNT concentration gradient and due to the tight spatio-temporal control of Wnt gene expression. Under these circumstances, lower levels of Wnt/β-catenin signaling are likely to occur and, as a result, part of the regulation may be due to the AXIN2 auto-inhibitory feedback loop. Therefore, it may be the ratio between the WNT and AXIN2 levels that is crucial to the regulatory role of AXIN2. We therefore used our model to explore the spectrum of possible β-catenin stabilizations under different WNT and AXIN2 levels. We ran a series of simulations with different initial WNT token levels: 3, 4 or 5, showing increased β-catenin stabilization in Fig 3A, and with different AXIN2 feedback strengths: the arc weight from t11 to AXIN was varied from 0 for no feedback to 0.15 for maximum feedback. As shown in Fig 6, we observed three different spectra of β-catenin stabilizations to the different initial WNT token levels. The highest β-catenin stabilizations (solid lines in Fig 6) were identical to those observed in Fig 3A (without AXIN2 feedback). At high feedback, the β-catenin stabilization is lowered, and a maximum appears after which the β-catenin level declines (dashed lines in fig 6). The lowest β-catenin stabilizations displayed three different peak responses. For the peak responses, the height of the peak and the duration of the response depended on initial WNT token levels. Maximal β-catenin stabilization comes later in the simulation for higher initial WNT token levels.

**Fig 6.**
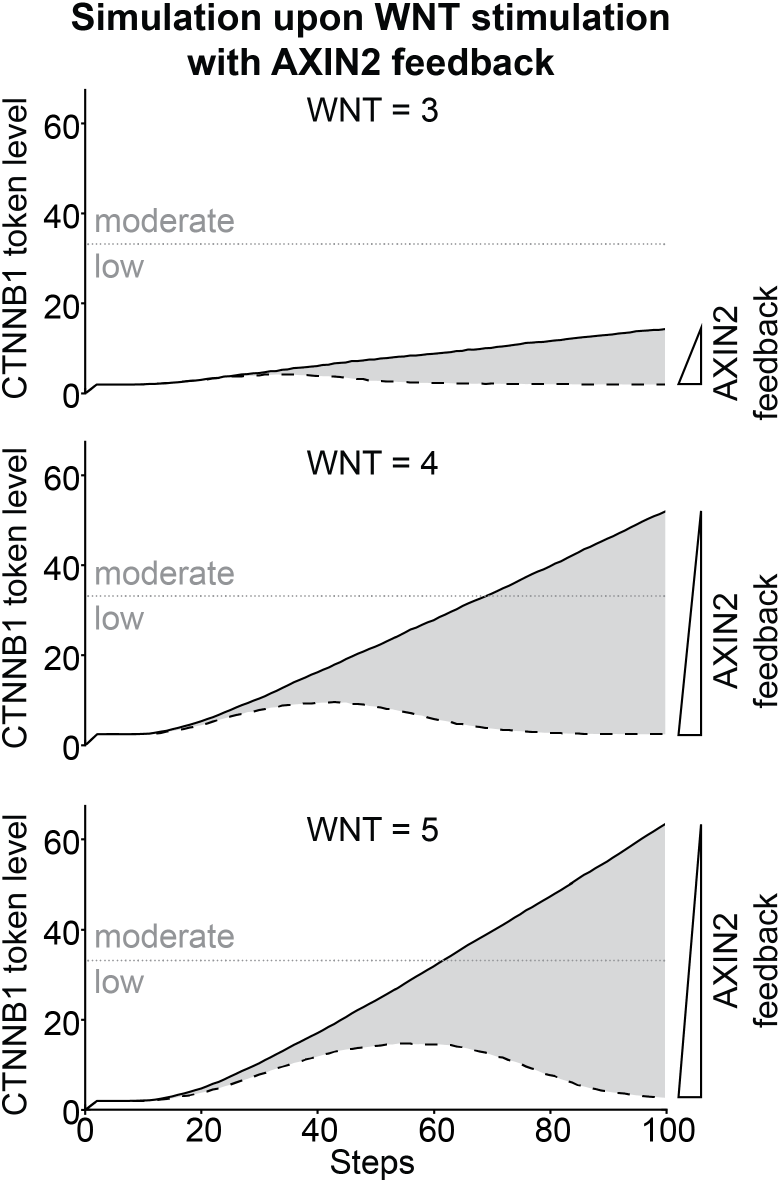
Model prediction of Wnt-pathway activation upon WNT addition with AXIN2 feedback. β-catenin (referred to by its official gene name CTNNB1 in the figure) token levels predicted by our model with arc weight from t11 to AXIN varied from 0 (no feedback; solid lines) to 0.15 (high feedback, dashed lines), and initial WNT token levels at 3, 4 and 5 (top, middle and bottom panels, respectively). We observed three spectra of β-catenin stabilizations depending on initial WNT levels. The highest β-catenin stabilizations correspond to simulations without AXIN2 feedback (solid lines), whereas with high AXIN2 feedback the β-catenin stabilization was attenuated (dashed lines).

## Discussion

In spite of more than 30 years of study, the Wnt/β-catenin signaling pathway still holds many questions. The molecular details of how an external WNT stimulus results in the stabilization of transcriptionally active β-catenin/TCF complexes remain incompletely understood and, in some cases, a topic of debate [41]. The Petri net model presented in this paper allows us to investigate downstream effects of Wnt/β-catenin signaling between different conditions. The core of the model describes the current state of knowledge as summarized in the introduction. AXIN1 was modeled as the only destruction complex component sequestered to the plasma membrane during Wnt/β-catenin signaling. An alternative mechanism, involving the actions of GSK3 at the plasma membrane, where it phosphorylates LRP5/6 has been suggested [42, 43], but we did not include this in the current model. Furthermore, we considered cytoplasmic and nuclear β-catenin as a single pool in the model. More detailed experimental data on subcellular compartmentalization of β-catenin (or any other signaling component) would allow us to refine our Petri net model, which easily allows incorporations of such detail.

Our model predicts a dose- and time dependent response for both WNT stimulation and GSK3 inhibition (Figs. 3A and 4A). This is confirmed by the experimental data (Figs. 3B, 3C, 4B and 4C). The main discrepancy between the simulated and the experimental data is the time-delay that is predicted in response to WNT stimulation compared to GSK3 inhibition (compare Fig 3A to 4A). Indeed, activation of Wnt/β-catenin signaling is known to be a slow event (unlike the activation of MAPK signaling for instance, which occurs within a matter of minutes) [44-46]. However, we did not detect this delay in β-catenin accumulation by either TCF/LEF luciferase reporter assay (compare Fig 3B-3D to Fig 4B-4D) or Western blot analysis (compare Fig 3E and 3F to Fig 4E and 4F). We believe this is mainly due to experimental limitations. Given that a subtle increase in β-catenin protein levels can be detected approximately one hour after stimulation with either Wnt3a (Fig 3E and 3F) or CHIR99021 (Fig 4E and 4F), any delay in activation of the Wnt-pathway must occur prior to that time point. Detecting this delay would require assays with superior spatio-temporal resolution. The delay predicted by our model upon WNT stimulation (Fig 3A) can be explained by the fact that formation of the signalosome occurs a few steps into the simulation, whereas inhibition of GSK3 is a one-step event, and that the transitions for signalosome formation (i.e. pathway activation) and destruction complex formation (i.e. pathway inhibition) compete for AXIN1. Thus, when AXIN1 is sequestered to the plasma membrane, less cytoplasmic AXIN1 is available for formation of the destruction complex. To what extent these events contribute to Wnt-pathway activation under experimental conditions remains unknown, owing to the absence of tools to study the exchange of AXIN1 between these two pools. Our results do suggest that competition over AXIN1 between the destruction complex and the signalosome may well be important also under physiological conditions.

The time-delay together with the continuous sequestration and dissociation of AXIN1 to the signalosome leads to prediction of higher stabilization of β-catenin for complete GSK3 inhibition compared to maximal WNT stimulation, where the difference is almost two-fold (compare Fig 4A to Fig 3A). We observe a similar difference when measuring TCF/LEF reporter gene activity: the highest concentration of CHIR99021 activates the reporter approximately 10-fold higher than the highest concentration of Wnt3a tested (compare Fig 4B-4D to Fig 3B-3D). Comparing protein levels, instead of transcriptional activation, shows a much smaller difference: 2-fold higher β-catenin at most when cells are stimulated with CHIR99021 versus Wnt3a (compare Fig 4E and 4F to Fig 3E and 3F). Although it is tempting to conclude that this data again confirms the predictions of our model, it should be stressed that the different experimental modes of Wnt-pathway activation cannot be compared directly. This is because they are achieved by different molecules (i.e. purified Wnt3a versus a synthetic small-molecule GSK3 inhibitor) with different intrinsic activities and chemical properties such as half-life and stability in the tissue culture medium, which may greatly impact on the experimental outcome. At the same time, we may speculate that the observed differences reflect real differences in sensitivity of the Wnt-pathway. In this case, our experimental findings might be explained by the fact that the more physiological means of pathway activation by Wnt3a is more likely to be subject to negative feedback control via AXIN2 induction than the more artificial perturbation by CHIR99021 inhibition of GSK3 at the level of the destruction complex.

AXIN2 is one of the few comprehensive globally expressed WNT target genes and is thought to act as a negative regulator to Wnt/β-catenin signaling [16, 17]. The degree to which AXIN2 attenuates WNT signaling and the actual spatio-temporal regulatory role of AXIN2 is still a topic of debate. Indeed, when we incorporate this negative feedback loop in the model upon WNT stimulation, our simulations predict that Wnt-pathway activity is attenuated (at certain levels of AXIN2 induction), and ultimately returns to baseline levels (dashed lines in Fig 6). Importantly, in the model the feedback from AXIN2 only negatively influences stabilized β-catenin levels when AXIN1 is the limiting factor. This is the case when AXIN1 is deprived from the cytoplasm by sequestration to the signalosome (i.e. upon WNT stimulation). This is why we are able to observe a negative effect from the AXIN2 feedback upon WNT stimulation in our model (fig 6), but not by GSK3 inhibition (Fig 4A) or APC inactivating mutations (Fig 5). It should be noted however, that on the timescale used for the experiments, we do not observe complete feedback inhibition by AXIN2 (Fig 3B-3F). This might be due to the relatively low level of *AXIN2* induction in the cells used for these experiments (data not shown) in combination with supra-physiological levels of Wnt-pathway activation achieved upon stimulation with purified Wnt3a. However, it could also be due to the fact that AXIN1 is not the limiting factor in the cells used for this study. Previously, a study of Wnt/β-catenin signaling in *Xenopus laevis* showed that AXIN1 is 1000-fold lower than the other components of the destruction complex [39] and has therefore been considered the natural limiting factor. However, a recent study of Wnt/β-catenin signaling in mammalian cells showed that the concentrations of the components of the destruction complex were on the same range [47]. Therefore, we cannot exclude the possibility that AXIN1 is not the limiting factor in the cells used for this study. Unfortunately, the current experimental tools, most notably Western blot analysis of endogenous β-catenin levels, are not sufficiently robust, high-throughput and sensitive enough to resolve this issue. However, by using our model we were able to predict and visualize spectra of β-catenin stabilization, which showed that the ratio between the WNT and AXIN2 levels are important for the degree of feedback observed (Fig 6). The two most notable observations were that, for high WNT levels, a higher level of AXIN2 was needed to reach baseline β-catenin levels and, for low WNT levels, a baseline β-catenin level is reached early. Based on these predictions we can speculate whether the AXIN2 negative feedback only has an effect on low WNT levels and whether the regulatory role of this is to insure a faster on/off switch of Wnt-pathway activity. Indeed, Wnt-pathway activity shows dynamic on and off switches during development [22]. Examples of these are the restriction of Wnt/β-catenin responsive cells to the crypt, but not to the villus sections of the intestinal epithelium, and oscillation of WNT signaling as part of the mouse segmentation clock.

In conclusion, our Petri net model of Wnt/β-catenin signaling provides insight on the mechanisms leading to different levels of β-catenin stabilization upon WNT stimulation and GSK3 inhibition corroborated by TCF/LEF luciferase assay and Western blot analysis. It should be stressed that the simulations show a coarse-grained output per step and we cannot directly map token levels to the relative activities in the TCF/LEF luciferase reporter assay nor to the β-catenin levels measured by Western blot analysis. Furthermore, we also cannot directly map a simulation step in the model to an experimental timescale. Despite these limitations, our model resembles Wnt/β-catenin signaling to the extent that it captures the logic of the interactions and reflects the sequence of events of pathway activation and repression by various mechanisms. In this way, our model can be used to simulate and predict both physiological and pathophysiological WNT signaling. Thus, this modelling exercise has allowed us to study the mechanisms and effects of Wnt/β-catenin signaling under different conditions, as well as the effects of protein- and pathway- modifications that are known to influence this pathway in many types of cancer.

## Materials and Methods

### Petri net modeling

We built a Petri net model of Wnt/β-catenin signaling describing known components, actions and interactions, well established in literature, in a logical way. A Petri net consists of two types of nodes, ‘places’ and ‘transitions’, and is connected by directed edges called ‘arcs’. A place represents an entity (e.g. gene or protein), whereas a transition indicates the activity occurring between the places (e.g. gene expression or complex formation). Places can only link to transitions and vice versa (i.e., a Petri net is a bipartite graph). The direction of the arcs is important for the flow of the network. An arc goes from an input place to a transition, and from a transition to an output place. Places contain ‘tokens’, indicating the availability of the corresponding entity, while arcs have a weight, denoting the amount of tokens to consume from an input place or to produce to an output place. If the token levels of all input places of a transition fulfill the requirement of (i.e. are equal to or higher than) the weights of the respective arcs, the transition is enabled. Only enabled transitions can be executed, leading to transfer (consumption/production) of tokens between places. Note that if two (or more) enabled transitions share an input place, they may be in competition if available token levels do not allow simultaneous execution of both (or all). In our model, AXIN, β-catenin and the destruction complex with β-catenin bound, are each input places for two transitions (t3/t5, t6/t10 and t7/t8, respectively).

Gene expression is modeled such that one arc goes from the gene-place to the transcriptional-transition, one arc goes from the transcriptional-transition to the gene-place, and one arc goes from the transcriptional-transition to the protein-place. When the transcriptional-transition of a gene is enabled a token is produced both in the protein-place and in the gene-place itself. This way the token can be reused for another round of gene expression, reflecting the fact that the gene (DNA) is needed, but is not consumed during expression.

### Active and hyperactive conditions in the model

We modeled active and hyperactive signaling upon WNT stimulation and GSK3 inhibition, respectively, and used these conditions to validate the model with experimental data (see below). Inhibition of GSK3 inhibits formation of the destruction complex, which we interpret to be similar to oncogenic perturbations. Therefore, for modeling purposes, GSK3 inhibition was used to mimic hyperactive signaling. For GSK3 inhibition we varied the initial token level of GSK3, respectively, from 0 to 5. For WNT stimulation we varied the initial token level of WNT from 0 to 5 and removed the AXIN2 feedback (the arc weight from t11 to AXIN was set to 0). The experimentally validated model was used to predict the level of β-catenin stabilization with the AXIN2 negative feedback upon WNT stimulation and APC inactivating mutations, respectively. Upon WNT stimulation with the AXIN2 feedback we varied the initial token level of WNT (3, 4 and 5) and the arc weight from t11 to AXIN (0 (no feedback) and 0.15 (maximal feedback)). Thus, the simulation for each initial WNT token level produced two β-catenin stabilization curves (i.e. no feedback and maximal feedback). The area between these two curves was used to explain the spectra of β-catenin stabilizations at intermediate levels of *AXIN2* induction. APC mutants have decreased binding affinity to the other components of the destruction complex to different degrees. We implemented this by reducing the formation of the destruction complex i.e. the weight on the arc going from the complex-formation-transition (t5) to the destruction complex (production) was decreased to 0, 0.05, 0.1 and 0.2. In addition, we incorporated arcs going from the complex-formation-transition to the individual destruction complex components (i.e. AXIN1, APC, GSK3 and CK1) with arc weights of 1 minus the production-weight to equally decrease the consumption. For a weight of 0 no destruction complex can be produced. For weights of 1/20, 1/10 and 1/5 the destruction complex was produced every 20, 10 and 5 steps, respectively.

### Simulations

The model was simulated with maximally parallel execution, cf. our previous work [37], where the maximum possible number of enabled transitions are executed at each simulation step. This mimics the behavior in the cell, where typically many interactions happen at the same time. Two or more transitions can compete over one input place, as mentioned above. If this place only contains enough tokens to enable one of the transitions, but not both, a conflict occurs which is resolved by randomly drawing one of the competing transitions to execute. This makes the simulations non-deterministic.

For each condition we simulated the total β-catenin token levels over 100 steps repeated 100 times. To account for variations in token levels due to the non-deterministic nature of the model, the mean and standard deviation of the β-catenin token levels over the 100 simulations were calculated for each step. The steps describe the sequence of events and should not be linearly translated to time units. Similarly, the token level is a coarse-grained quantitative representation of actual protein levels and should not be linearly translated to a concentration. Instead, for analysis of the simulations we observe relative differences of β-catenin token levels over steps between simulations (i.e. different conditions and dosages). To validate the model we compared the β-catenin levels predicted by the model simulations to the Wnt-pathway activities measured in experiment (see below). A Python script was written to run the simulations and is available together with the model in pnml format via http://www.ibi.vu.nl/downloads/WNTmodel/.

### Cell lines

HEK293T^WOO^ (WNT OFF/ON) cells were generated by transfecting HEK293T cells with a 7xTcf-FFluc//SV40-Puro^R^ (7TFP) reporter plasmid (a gift from Christophe Fuerer, [48]). Following puromycin selection to obtain stable integrants, individual clones were assessed for their response to Wnt-pathway activation. The clone with the highest dynamic range was used for the experiments depicted in Figs. 3 and 4.

### Cell culture and stimulation

HEK293T^WOO^ cells were cultured in Dulbecco’s Modified Eagle Medium: Nutrient Mixture F-12 (DMEM/F12) supplemented with 10% FCS and 1% Penicillin/Streptomycin (GIBCO, Life Technologies) in 5% CO_2_ at 37 °C. These cells respond to activation of the Wnt/β-catenin signaling pathway by expressing firefly luciferase, since the firefly luciferase in the 7TFP construct is driven by the 7xTcf promoter, which contains 7 repeats of the TCF/LEF transcription response element. Cells were plated the day prior to stimulation in a 96 well-plate at a density of 20.000 cells per well. Cells were stimulated with different concentrations (10-200 ng/ml) of purified Wnt3a protein (RnD) dissolved in 0.1% BSA in PBS, or with different concentrations (750 nM-6 µM) CHIR99021 (BioVision) dissolved in DMSO, for different amounts of time (1-24 hours). At the indicated time points following stimulation, cells were lysed in 20 µl of Passive Lysis Buffer (Promega) and cell lysate from the same experiment was used for both the luciferase assay (3 wells per condition) and Western blot analysis (the remainder of the 3 wells, pooled per lane).

### Western blot analysis

Protein concentration was measured using the Pierce BCA Protein Assay Kit (Thermo Fisher Scientific). Equal amounts of protein were run on an 8% SDS-PAGE gel. Proteins were transferred to a nitrocellulose membrane (Bio-Rad) and blocked with TBS Odyssey Blocking Buffer (LI-COR Biosciences, diluted 1:1 in TBS prior to use). Primary antibodies directed against active β-catenin (Cat# 8814S, Cell Signaling, 1:1000), total β-catenin (Cat# 610153, BD Biosciences, 1:2000) and α-Tubulin (Cat# T9026, Sigma-Aldrich, 1:500) were diluted in blocking buffer supplemented with 0.1% Tween-20 (TBS-T). Staining was performed overnight at 4 °C. Membranes were washed in TBS-T followed by incubation with secondary antibodies (IRDye 680LT (Cat# 926-68021) or IRDye 800CW (Cat# 926-32212) (LI-COR), 1:20000 in TBS-T) for 2 hours. Membranes were washed in TBS-T and incubated in TBS prior to scanning at 700 nm and 800 nm using an Odyssey Fc (LI-COR Biosciences). Image Studio^TM^ Lite 4.0 software (LI-COR Biosciences) was used to quantify relative protein levels. Background correction was performed according to the manufacturer’s instructions (median of pixels, top/bottom border width of 3).

### Luciferase assay

To measure the activity of firefly luciferase (and hence Wnt-pathway activity), 10 μl of cell lysate was transferred to a black 96-well Optiplate (Perkin Elmer). The SpectraMax L Microplate Luminometer was used to inject 50 μl Luciferase Assay Reagent II (Promega) per well followed by measurement of firefly luciferase activity.

## Acknowledgment

We thank Vivienne Woo for helping to establish and characterize the HEK293T 7TCF reporter cell line.

